# Large-scale structural analysis of pre-mRNA structure using RADIS

**DOI:** 10.64898/2026.09.08.750205

**Authors:** Claire Fleurisson, Sijin Chen, Ryan M. Macdonald, George S. Stephenson, Hao Wu, Anastacia N Wienecke, Saumya Gupta, Peter J. Castaldi, Ayan Paul, Kevin M. Weeks, Alain Laederach

## Abstract

Pre-mRNA secondary structure can modulate the regulatory function of intronic sequences by masking or exposing splice-site signals and altering the accessibility of other features. Yet experimentally supported structural models for human pre-mRNAs have been largely restricted to splice-site-proximal sequences, leaving the deep intronic regions where ∼70% of intronic disease-causing variants reside almost entirely unmapped. We developed RADIS (Reactivity Analysis of Deep and Intergenic RNA Structure), a strategy that yields strand-resolved chemical probing reactivity profiles across entire intron-rich loci by creating tiled arrays of RNAs that comprehensively span long genomic distances. This approach removes the abundance and locus-specificity constraints that otherwise limit pre-mRNA probing approaches. RADIS recovers known *E. coli* ribosomal RNA architectures and yields high-correlation with in-cell dimethyl sulfate (DMS)-MaP reactivities at representative splice sites (*r* = 0.87–0.90). RADIS-constrained folding of 109 5′ and 88 3′ splice sites uncovers an inverse relationship between base-pairing within the spliceosome footprint and splice-site strength, and partitions 81 branchpoints into three structural classes. Across 233 full-length Alu elements, sense Alu RNAs are more structured than antisense elements, and both partition into strand- and lineage-dependent structural classes. RADIS enables experimentally grounded structure analysis at disease-associated intronic and intergenic loci, complementing sequence-based variant-effect predictors.

## Introduction

RNA structures at and near splice sites directly influence splice site recognition by masking binding sites for the spliceosomal machinery or bringing distant splice sites into proximity. For example, RNA hairpins at 5′ splice sites have been found to occlude U1 snRNP binding, directly suppressing splice site recognition (Warf and Berglund 2010; Rubtsov 2016; Kumar et al. 2022; Cao et al. 2024). Structure at branch points and 3′ splice sites can similarly gate their recognition (Buratti and Baralle 2004). Despite this mechanistic importance, obtaining accurate, experimentally supported structure models for endogenous pre-mRNAs remains an outstanding challenge, especially for deep intronic regions. Introns make up ∼88% of transcribed pre-mRNA sequence, and ∼11% (over 25,000) of known disease-causing mutations map to introns. Of these disease-causing mutations, ∼70% map outside of 5′ and 3′ splice sites, illustrating the importance of deep intronic sequences (Waldern et al. 2022; Waldern et al. 2025).

Pre-mRNAs are present at far lower abundance than mature mRNA, are rapidly processed, and their intronic sequences are poorly captured by transcriptome-wide chemical probing approaches. Methods such as DMS-MaP (Mustoe et al. 2019) have advanced transcriptome-wide structural studies, yet sensitivity for low-abundance intronic regions remains limiting (Bubenik et al. 2021; Rangan et al. 2022; Eich et al. 2024; Yarshova et al. 2026). Gene-specific RT-PCR with an exon-anchored primer enabled selective amplification of unspliced intronic sequence for DMS-MaP probing at splice-site-proximal regions (Kumar et al. 2022). However, this strategy requires relatively high endogenous pre-mRNA levels and specific primer design for each locus. For disease-relevant genes expressed at low levels in specific cell types, such as *ESR1* [associated with breast cancer (Baek et al. 2025; Li et al. 2025; Blanc-Durand et al. 2026)], *BRCA1* [associated with breast/ovarian cancer and containing multiple Alu-rich introns (Auranen et al. 2005; Mucaki et al. 2016)], and *DENND1A* [associated with polycystic ovary syndrome (PCOS) (Tee et al. 2016)], these limitations effectively preclude systematic structural profiling of pre-mRNA at a gene-wide scale.

Direct experimental measurement of RNA structure has been transformed by high-throughput chemical probing. Selective 2′-hydroxyl acylation (SHAPE) and dimethyl sulfate (DMS) probing detect conformationally flexible, often single-stranded, nucleotides. Mutational profiling readouts such as SHAPE-MaP and DMS-MaP encode each modification as a cDNA mismatch, allowing per-nucleotide reactivities to be quantified at scale (Siegfried et al. 2014; Mauger et al. 2015; Smola et al. 2015; Zubradt et al. 2017; Boerneke et al. 2023). Applied transcriptome-wide, these chemistries have revealed pervasive, regulation-relevant structure in bacterial (Mustoe et al. 2018), yeast and human transcriptomes (Ding et al. 2014; Rouskin et al. 2014; Wan et al. 2014; Spitale et al. 2015), and in viral genomes (Mauger et al. 2015; Dethoff et al. 2018; Kutchko et al. 2018; Tomezsko et al. 2020; Allen et al. 2025). However, these approaches sample the steady-state RNA pool, which is dominated by mature messages. pre-mRNAs and their intronic sequences are typically two to three orders of magnitude less abundant and are further depleted by polyA selection in most library protocols. Compounding these challenges, co-transcriptional folding couples splicing decisions to nascent RNA structures that can be transient on the timescale of typical chemical-probing reactions (Saldi et al. 2016; Watters et al. 2016; Saldi et al. 2021). The structural features most relevant to splice-site recognition (5′-splice-site hairpins, branchpoint context, and deep-intronic regulatory motifs) have thus remained difficult to probe at endogenous loci and have most often been inferred from short synthetic RNAs studied outside their genomic context (Eperon et al. 1988; Buratti and Baralle 2004).

The therapeutic and diagnostic importance of understanding pre-mRNA structure is well established. Single-nucleotide variants that perturb secondary structure (riboSNitches) account for a disproportionate fraction of disease-associated non-coding alleles and can rewire splicing, miRNA accessibility, and protein binding (Halvorsen et al. 2010; Kutchko et al. 2015). Translation of this principle into the clinic has been most dramatic in spinal muscular atrophy, where structural insights at the SMN2 intron-7/exon-7 junction enabled both antisense-oligonucleotide (Hua et al. 2008) and small-molecule splice-modulating therapies (Naryshkin et al. 2014). Analogous structure-aware strategies are being explored for cancer-associated splicing dysregulation (Bonnal et al. 2020). Despite the maturity of chemical-probing technologies, the field has lacked a strategy for obtaining robust, gene-wide pre-mRNA structural profiles for such loci independent of their endogenous transcript abundance or of the cell type in which they are expressed. RADIS (Reactivity Analysis of Deep Intergenic RNA Structure), described here, fills this gap by decoupling RNA structural interrogation from in vivo transcription. Any locus carried on a (large) plasmid, including bacterial artificial chromosomes (BACs), can be transcribed, probed, and structurally annotated efficiently. RADIS provides an experimentally grounded structural map onto which variant effects and RNA structure-driven processing aberrations can subsequently be projected. While RADIS is demonstrated here on pre-mRNA loci, the strategy is broadly applicable to intronic long non-coding RNAs and to any genomic region that can be cloned into a plasmid or BAC.

Here, we validate the approach using highly structured bacterial rRNAs as an internal benchmark, demonstrate that RADIS reactivity profiles correlate with in-cell DMS data at splice-site-proximal regions, and apply RADIS to BACs covering *ESR1*, *BRCA1*, and the *CRB2*–*DENND1A* locus to obtain new insights into splicing and Alu element structure. RADIS ultimately provides a scalable framework to map pre-mRNA secondary structure across full genomic loci.

## Results

### Workflow for strand-resolved DMS reactivities across complete pre-mRNAs

To enable RNA structure probing of pre-mRNAs, we integrated the RADIS workflow with bacterial artificial chromosome (BAC) DNA to yield strand-resolved DMS reactivity profiles suitable for secondary structure modeling (Figure 1A). BACs carrying human genomic inserts (Figure 1B) were fragmented by ultrasonication, empirically optimized to generate fragments above 1 kb. This fragment length was chosen as a compromise between preserving sufficient long-range structural information and remaining compatible with efficient library preparation and sequencing. Fragments showed a unimodal size distribution with a mean of 1.5 kb, above the 1 kb target, and with most fragments falling between 0.7 and 3.0 kb (Supplementary Figure 1A).

**Figure 1:**
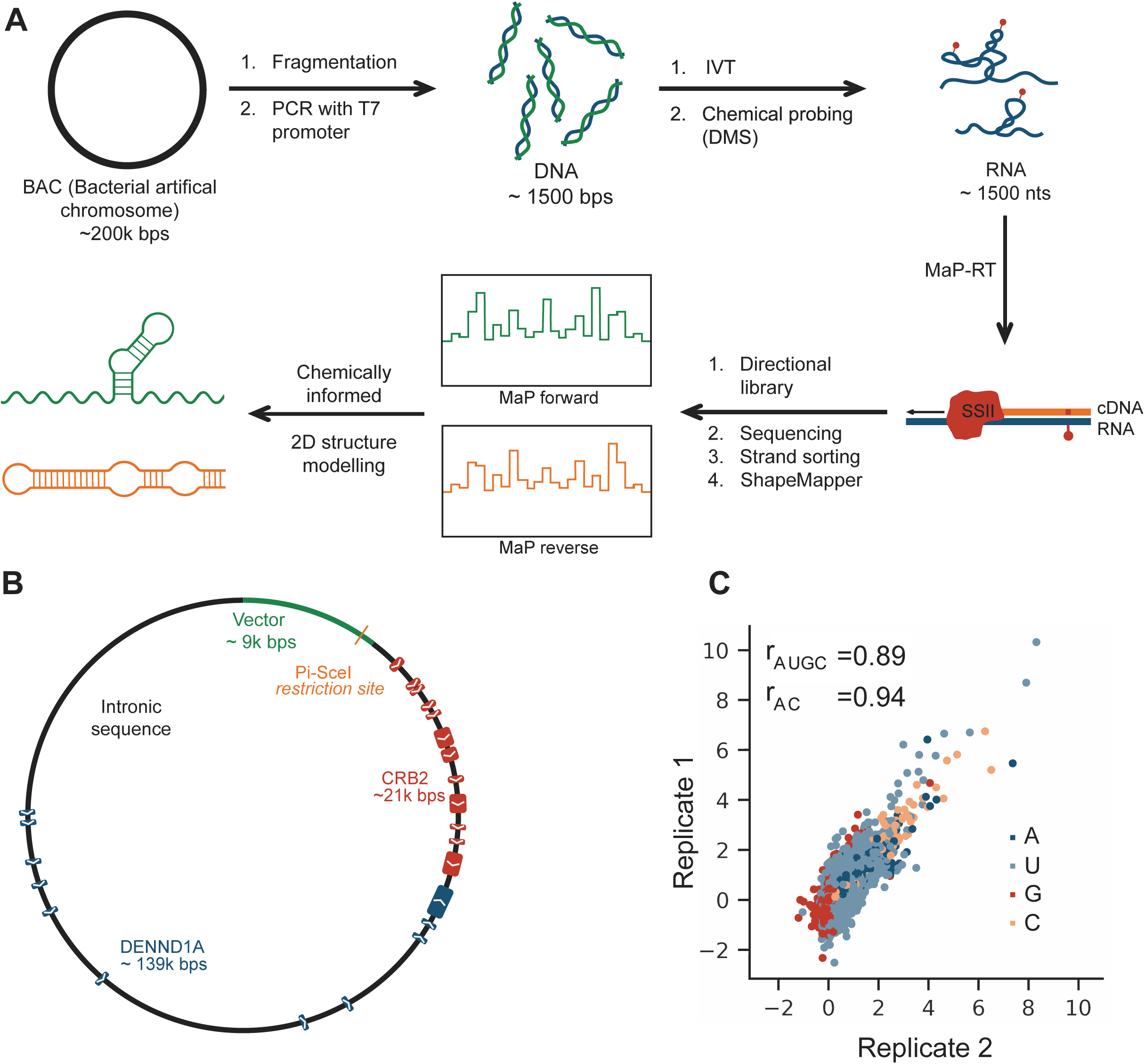
RADIS strategy. (A) Overview of the RADIS workflow, including: BAC DNA fragmentation and *in vitro* transcription, RNA chemical probing, MaP-RT, and secondary structure modeling. (B) The RP11-147E3 BAC covering chr9:123357475–123518426 (not to scale). (C) Correlation of per-nucleotide DMS reactivity profiles for independent RADIS replicates. Pearson correlation coefficients are shown for A/C-only and all nucleotides.

To append a T7 promoter to fragmented BAC DNA, primers were redesigned so that the (Illumina platform) i7 adaptor carried a T7 promoter sequence at its 5′ end, and then prepared templates using a commercial library preparation approach (NEBNext Ultra II DNA). The resulting libraries serve directly as templates for *in vitro* transcription. High-complexity DNA libraries were thus obtained without the need for additional ligation steps, and *in vitro* transcription with T7 RNA polymerase then generated RNA corresponding to both strands of the DNA (BAC) fragments.

DMS probing was carried out using established bicine-buffered conditions, followed by MaP-RT to encode modification events as mutational signatures (Krokhotin et al. 2017; Mustoe et al. 2019; Grayeski et al. 2022). Because transcription occurs from both strands, this approach generates both sense and antisense RNAs. We retained strand information using directional cDNA library generation after MaP-RT. This step was critical for unambiguously assigning reactivities to the correct genomic strand in regions with overlapping transcription.

### Reproducible DMS data across entire pre-mRNA of disease genes

Sequencing was carried out using the Illumina platform, and median per-nucleotide read depth ranged from 2545 to 13053 (Supplementary Table 1). Reads were aligned to the BAC insert sequence using a strand-aware mapping strategy, and alignments were sorted by strand to reconstruct separate reactivity profiles for sense and antisense transcripts (Wu et al. 2026). Secondary structures were predicted by incorporating DMS restraints (Mustoe et al. 2019) as implemented by EternaFold (Wayment-Steele et al. 2022), which yielded substantially better agreement with known structures than conventional thermodynamic folding methods, producing base-pairing models for selected regions and for specific structural elements of interest.

RADIS was applied to three BACs selected for their primary gene content and disease relevance. One BAC harbored *ESR1*, a key breast cancer gene (Baek et al. 2025; Li et al. 2025; Blanc-Durand et al. 2026). The second contained the *CRB2*–*DENND1A* locus (Figure 1B), where *DENND1A* is implicated in polycystic ovary syndrome. The alternatively spliced isoform *DENND1A*.V2 is overexpressed in PCOS theca cells and sufficient to confer a PCOS androgen-overproduction phenotype (Tee et al. 2016). A third *BRCA1*, which is associated with breast cancer and enriched in Alu elements (Auranen et al. 2005; Mucaki et al. 2016). DMS reactivity profiles were highly reproducible across all probed genes. For example, the *CRB2*–*DENND1A* sense-strand RNA showed Pearson correlation coefficients of 0.89 when calculated over all nucleotides and 0.94 when restricted to A/C positions (Figure 1C). Gene-wise Pearson correlation coefficients between replicates ranged from 0.75 to 0.93 for all nucleotides and from 0.93 to 0.97 for A/C positions, and global median reactivity values fell within a narrow range (Table 1). Together, these results indicate that the BAC-based RADIS workflow yields consistent, strand-resolved DMS reactivity profiles over RNAs spanning ∼170 kb.

**Table 1.** RADIS analysis of target genes. BAC size, probed genes, replicate reproducibility, and median nucleotide reactivity are shown. Replicability values are Pearson correlation coefficients between independent replicates calculated over all nucleotides (AUGC) and over A/C positions only. Median reactivity values are the median [25th–75th percentile] of per-nucleotide mean reactivities across two biological replicates for each gene. Asterisks indicate genes that are only partially contained within the BAC.

| BAC | Size<br>(nts, kb) | Gene | Replicability<br>AUGC, AC | Median reactivity |
| --- | --- | --- | --- | --- |
| RP11-95G8 | 169 | *ESR1\** | 0.75, 0.93 | 0.39 [0.19, 0.63] |
| RP11-147E3 | 161 | *CRB2\** | 0.92, 0.95 | 0.34 [0.16, 0.60] |
|  |  | *DENND1A\** | 0.89, 0.94 | 0.40 [0.23, 0.64] |
| RP11-812O5 | 163 | *BRCA1* | 0.91, 0.97 | 0.41 [0.23, 0.64] |
|  |  | *NBR2* | 0.93, 0.97 | 0.36 [0.17, 0.61] |
|  |  | *NBR1\** | 0.92, 0.97 | 0.39 [0.19, 0.63] |

### Quantitative validation of DMS reactivities obtained on known RNA structures

We validated RADIS using an internal reference derived from the *E. coli* host. During BAC DNA preparation, a small fraction of chromosomal *E. coli* DNA co-purifies with the BAC, including through *in vitro* transcription and DMS probing. *E. coli* RNA fragments (∼1.5 kb) are processed by the RADIS pipeline in parallel with BAC-derived transcripts. We concatenated reads from all RADIS experiments and extracted DMS reactivity profiles for the 16S and 23S rRNAs. These profiles were compared with an independent cell-free DMS-MaP probing dataset acquired under similar bicine-buffered conditions (Mitchell et al. 2023).

RADIS and cell-free reactivity profiles were strongly correlated across ribosomal RNAs. For 16S rRNA, per-nucleotide reactivities yielded Pearson correlation coefficients of 0.81 over all nucleotides and 0.87 when restricted to A/C positions (Figure 2A); for 23S rRNA, the corresponding values were 0.86 and 0.87 (Figure 2B). Visual inspection revealed predominantly punctate differences, with slightly elevated RADIS reactivities at a subset of positions, consistent with the loss or weakening of certain long-range interactions upon fragmentation. DMS-constrained modeling of these profiles recovered the accepted 16S and 23S architectures with comparable accuracy from RADIS and cell-free data (16S rRNA, AUC = 0.74 for RADIS versus 0.79 for cell-free; 23S rRNA, 0.75 versus 0.78; Supplementary Figure 1B and 1C).

**Figure 2:**
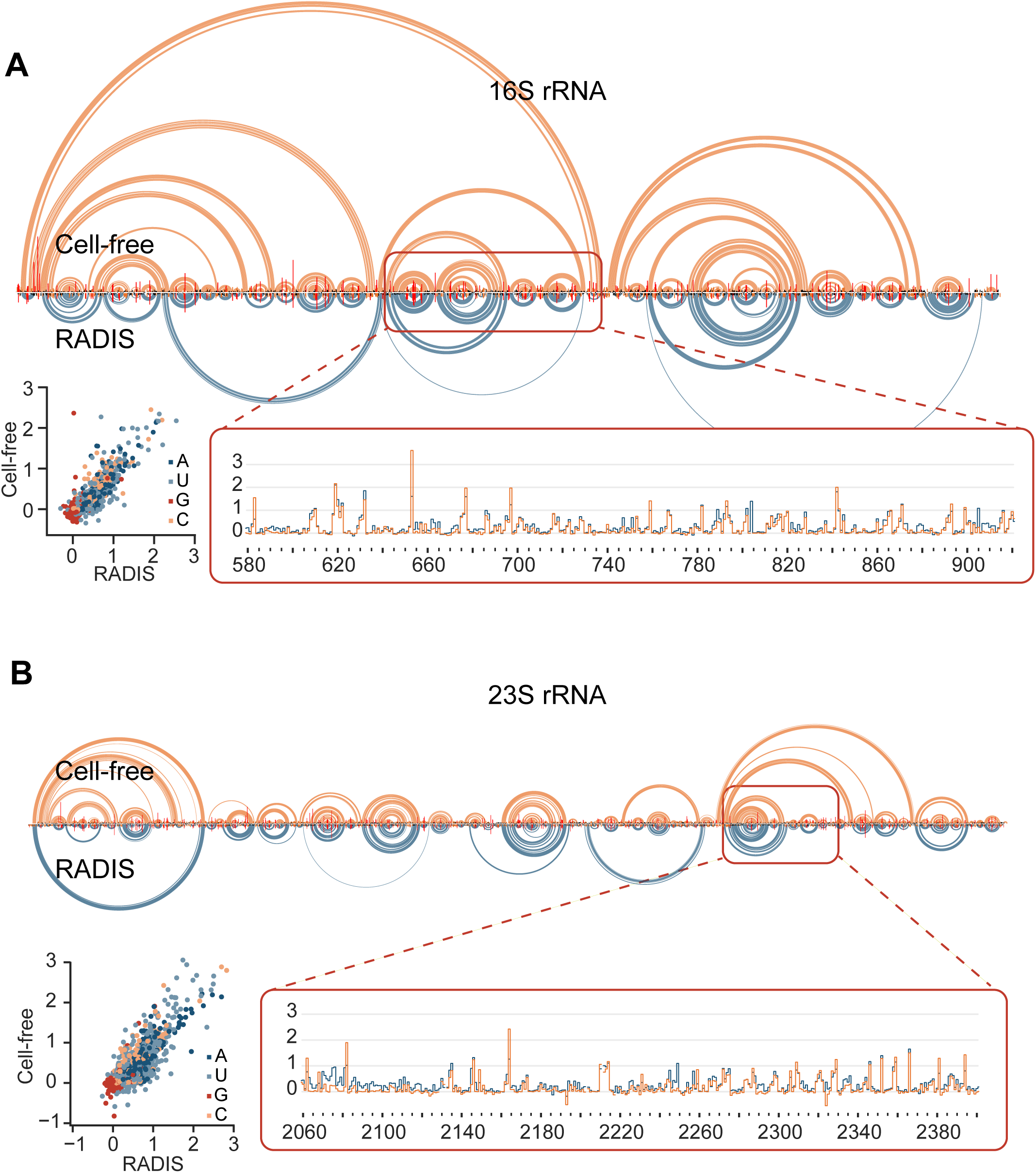
Comparison of RADIS and cell-free DMS probing of *E. coli* rRNA. (A) Arc plots of secondary structure models of 16S rRNA based on DMS reactivity-constrained modeling using native, cell-free RNA (orange) (Mitchell et al. 2023) and RADIS (blue). Expanded view of per-nucleotide reactivities for a representative region is shown at lower right. Scatter plot shows per-nucleotide reactivity correlation between cell-free and RADIS measurements (r_AUGC_ = 0.81 and r_AC_ = 0.87). (B) Comparison of cell-free and RADIS probing for 23S rRNA (r_AUGC_ = 0.86 and r_AC_ = 0.87).

We modeled secondary structures for both datasets using UltraFold, a windowed implementation of the SuperFold approach (Smola et al. 2015), including preservation of the 5′ loop and a loop near the 3′ end. When RADIS reactivities were evaluated against the R-scape base-pairing annotation for the microRNA MIR601 encoded antisense to DENND1A, the resulting ROC analysis yielded an AUC of 0.67. This indicates that the MIR601 RADIS profile carries structural information (Supplementary Figure 2A). This value is modest relative to the rRNA benchmarks, likely reflecting both the sparser, covariation-derived reference available for this RNA and the possibility that the in vitro-transcribed conformation differs modestly from it.

Representative RADIS data from *CRB2* intron 9 were then used to examine local structure around individual splice junctions. The RADIS-constrained secondary structure revealed distinct base-pairing patterns at representative 5′ and 3′ splice sites (5′SS and 3′SS), including both base pairs entirely within the intron or exon and base pairs that bridge exon–intron boundaries (Supplementary Figure 2B). These examples illustrate that RADIS can capture structured elements directly overlapping annotated splice sites and branchpoint regions within a long intron.

### Splice-site and branchpoint structures validated in cells

To assess and compare RADIS-derived structures relative to in-cell behavior, DMS probing was also performed in living cells for a subset of splice sites. Three splice sites in *ESR1* and four splice sites in *NBR1* were probed in cells and the resulting in-cell DMS profiles were normalized by boxplot transformation. We illustrate these comparisons for two splice sites of the *ESR1* gene (Figure 3A, 3B). For both the 5′SS and 3′SS, Pearson correlation coefficients between in-cell and RADIS reactivities are 0.87 to 0.90 (Figure 3C).

**Figure 3:**
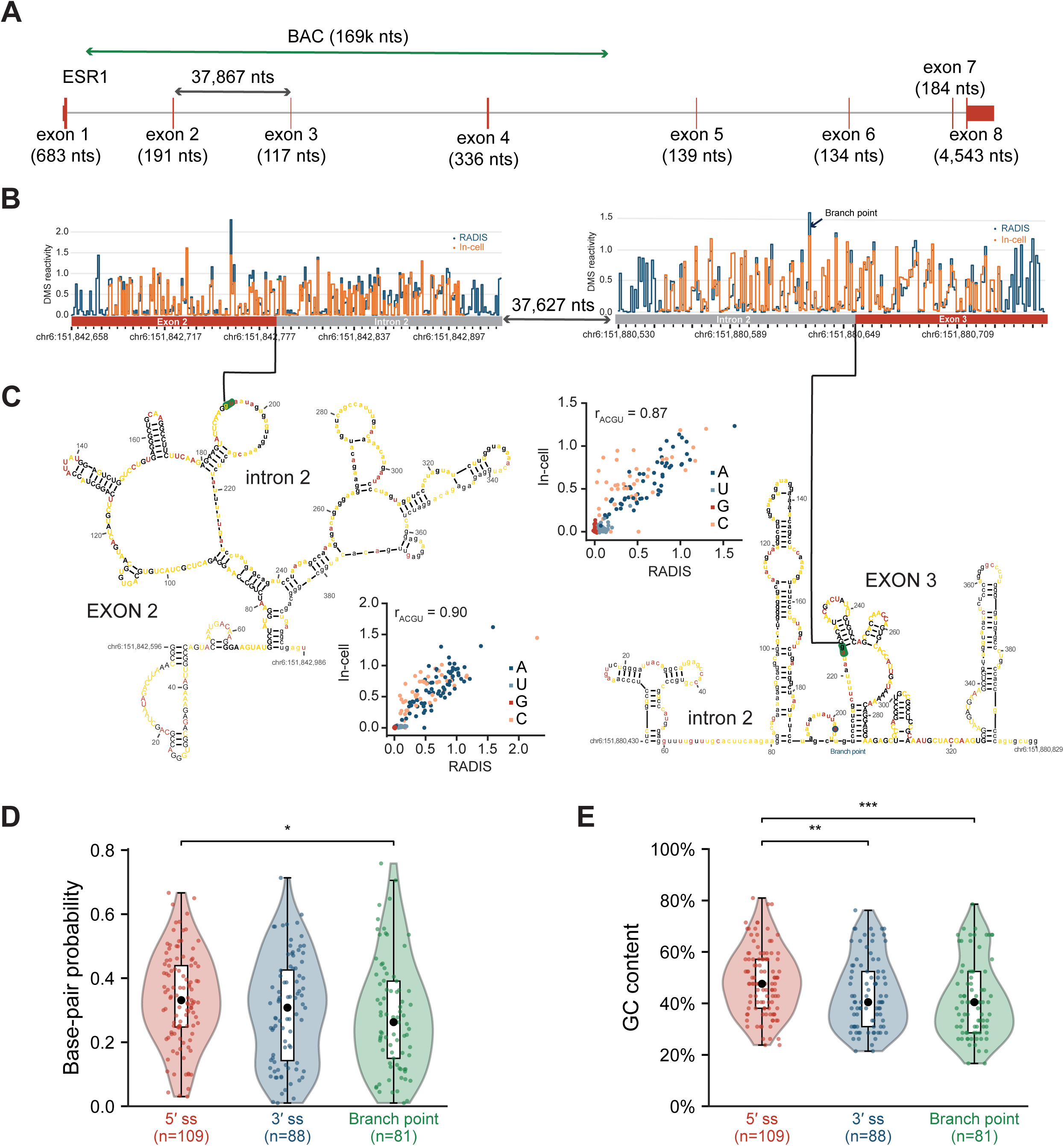
RADIS data are highly correlated with in-cell DMS probing at 5′ and 3′ splice sites and reveal complex pre-mRNA structures. (A) Architecture of the *ESR1* gene, containing 8 exons. The RP11-95G8 BAC spans 169 kb, and spans introns 1-4. (B) Comparison of in-cell DMS structure probing (orange) and RADIS data (blue) at the 5′ and 3′ splice sites that define intron 2. (C) Intron junction structure and correlation between RADIS and in-cell DMS probing. r_ACGU = 0.87 to 0.9. RADIS-constrained secondary structure models are shown for a representative EXON–intron junction-spanning region, with exon sequence in capital letters and intron sequence in lowercase letters; the region effectively probed in-cell is shown in bold. The 2D structure model spans the ∼400 nt surrounding the splice-site footprint and was used in subsequent analyses. Similar quantitative agreement was observed for five additional splice sites (Supplementary Figures 3 and 4). (D) DMS-informed base-pair probabilities for 5′ splice sites are higher than those for branchpoints, consistent with (E) average GC content stratified by site class.

High correlations were similarly observed for the five additional splice sites in *ESR1* and *NBR1* that were probed in cells (Supplementary Figures 3 and 4). These data show that the RADIS approach largely recapitulates in-cell DMS reactivity. When we used these data to inform RNA secondary structure models of these splice sites, we observed complex structures (Figure 3C). Exonic, intronic, and junction-spanning structures are modeled, indicating significant potential for RNA structure at these splice sites, and likely splice sites in general.

**Figure 4:**
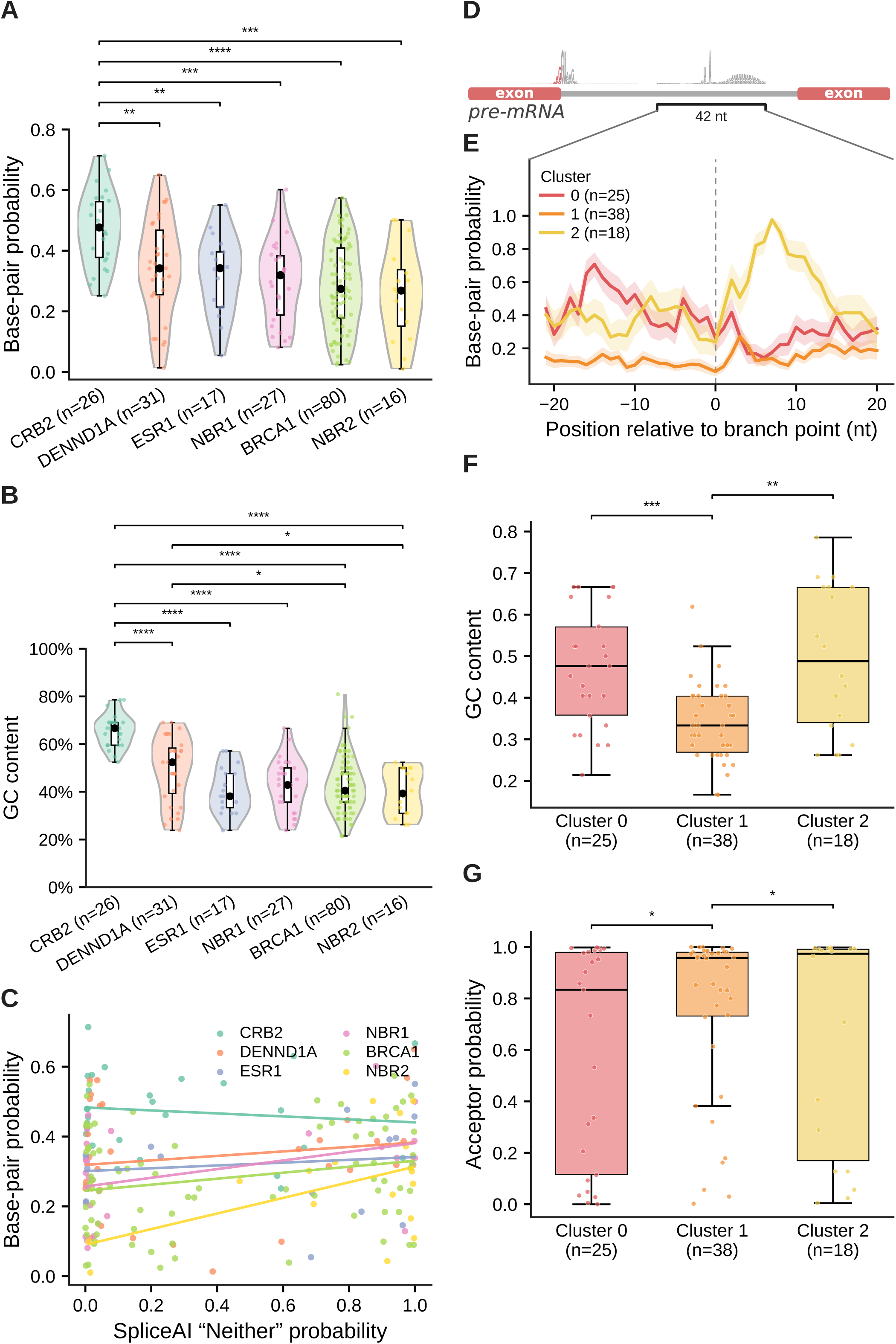
Summary of structural features measured by RADIS. (A) Base-pairing probabilities across pre-mRNAs (*CRB2*, *DENND1A*, *ESR1*, *NBR1*, *BRCA1*, and *NBR2*). Pairing probabilities are shown as violin plots; boxplots indicate the median (center line) and interquartile range (IQR); the mean is shown as a dot. Significance by two-sided Mann-Whitney U test. (B) Violin plots of GC content across the same pre-mRNAs. Statistical significance assessed using two-sided Mann-Whitney U tests. (C) Relationship between SpliceAI “neither” probability and average base-pairing probability at splice sites (3′ and 5′ splice sites), with points colored by gene identity. Lines indicate RNA-specific trends in structure-splicing relationships. (D) Scheme for analyzing the 42-nucleotide spliceosome footprint, centered at the branch point. (E) Mean base-pairing probability centered across branch points (n = 81), stratified by cluster (Clusters 0–2). Mean ± SEM is shown with shading. (F) GC content for each branch point region cluster. Boxplots show the median and 25th and 75th percentiles (box limits); whiskers extend to 1.5× the interquartile range (IQR). (G) SpliceAI acceptor probability of the 3′ splice site associated with each branchpoint, across branch point clusters, shown as box plots. For all plots, *, p < 0.05; **, p < 0.01; ***, p < 0.001; ****, p < 0.0001; two-sided Mann-Whitney U test. p-values in (A) and (B) are Benjamini-Hochberg FDR-adjusted for multiple comparisons; those in (F) and (G) are uncorrected. In (G), acceptor probabilities ≥ 0.98 were excluded from the test.

### Global analysis of splice sites in CRB2, DENND1A, ESR1, NBR1, BRCA1, NBR2

We examined a 400 nt folding window-around each splice site to capture the relevant local structural context while maintaining a tractable base pairing search space. This choice was based on an analysis of rRNA folding where we found that AUC values for 300-nucleotide windows were lower than for windows of 400–600 nt, which produced similar mean and median AUC and Pearson correlation values (Supplementary Table 2).

The spliceosome interacts with a contiguous pre-mRNA region that must be unfolded to be engaged, which we define as the spliceosome footprint. Based on analysis of cryo-EM structures across the spliceosome assembly pathway, we previously found that this footprint is largest in the catalytically activated B^act^ complex, where an approximately 42-nucleotide region spanning the 5′SS must become single-stranded. The unfolding energy of this region is the strongest structural predictor of splicing outcome (Kumar et al. 2022). We similarly found the 3′ splice-site footprint to comprise 42 nucleotides, 12 exonic and 30 intronic. Because 3′ splice-site usage is initiated at the branchpoint, we extended this concept by measuring, in the same activated (B^act^) spliceosome structure, PDB 5Z56 (Zhang et al. 2018), the pre-mRNA segment enclosed around the branch site, which is likewise approximately 42 nucleotides and which we used as the branchpoint footprint in following sections of this work. Unfolding of this branchpoint window is required for branch-helix formation and is completed during the Prp2-driven B^act^-to-B* transition that catalytically activates the spliceosome. Apparently, the spliceosome consistently interacts with roughly 42-nt long sequences at the 5′SS, 3′SS and branchpoint.

For each 5′SS, 3′SS and branchpoint, a 400 nt sequence-centered window was folded using RADIS DMS reactivities as constraints. Base pairing probabilities were extracted for the 42-nucleotide spliceosome footprint corresponding to each motif. Base-pair probabilities were averaged across each footprint to obtain a mean for each site. Comparison of these site-wise means and medians revealed significant differences among splice-site classes (Kruskal–Wallis test followed by pairwise two-sided Mann–Whitney U tests; p < 0.05)) (Figure 3D). 5′ splice-site footprints displayed higher average base-pair probabilities (median 0.34) than branchpoint footprints (median 0.26), with 3′SS showing intermediate values. Base pairing probabilities are correlated with GC content (Figure 3E) as expected (Signal et al. 2018; Monger et al. 2019). These analyses indicate that RADIS can be used to quantify and compare local structural propensities around large numbers of splice sites within BAC-derived transcripts and that these models detect differences among splice-regulatory regions.

### Analysis of structure/function in alternative splicing

GC content varies across the genome, which influences average spliceosome footprint base-pair probability on a gene-by-gene basis (Figure 4A). *CRB2* has the highest overall GC content (Figure 4B) and the highest base-pair probability in our RADIS-informed structural models; *NBR2* has the lowest values for both. Base-pair probability of spliceosome footprints is correlated with GC content, as expected. We next asked whether footprint structure, beyond its expected dependence on GC content, influences alternative splice-site usage.

We identified the alternative, constitutive and cryptic splice sites across the six pre-mRNAs in our RADIS data using the HEXEvent database (Busch and Hertel 2013). We obtained 109 5′SS and 88 3′SS. Branchpoints were assigned for 81 introns using published branchpoint maps and Branchpointer predictions (Signal et al. 2018). For each site, a 42-nt structural footprint was defined, as described above: 12 and 30 nucleotides in the exon and intron, respectively, for each 5′SS and 3′SS. A 42-nucleotide centered window was used for each branchpoint. We predicted the SpliceAI (Jaganathan et al. 2019) acceptor or donor probability and found that these scores correlated strongly with the fraction of transcripts in which each site is constitutively used, as catalogued in HEXEvent (Supplementary Figures 5A and B). We used SpliceAI for subsequent analyses because it provides genome-wide predictions, more comprehensive than the tissue-specific HEXEvent catalogue (Jaganathan et al. 2019), and could be computed for every splice site and branchpoint across our data.

We first examined the “neither probability” from SpliceAI (Figure 4C). The neither probability is a measure of splice site strength, with 0- and 1-values indicating strong and weak splice sites, respectively. This score reflects the probability that a position is a splice site rather than the extent to which it is used. We used this metric rather than the separate donor and acceptor probabilities because these analyses treat donors and acceptors together, and the probability that a nucleotide is neither a donor nor an acceptor applies equally to both. We see the strongest correlation between neither probability and base-pair probability in the *NBR2* gene, which has the lowest level of structure and GC content. Conversely, we observe a weak negative correlation for *CRB2*, the gene with the highest GC content. Thus, the presence of a strong (highly paired) structure in the spliceosome footprint region has a larger negative effect on splice site strength in genes with lower GC content. This observation is consistent with the idea that the spliceosome preferentially binds to relatively unstructured regions of a pre-mRNA, making low structure regions stronger splice sites.

### RADIS analysis of branchpoint structures

The data we have presented so far considers the mean and median base-pairing probability of a (42-nt) spliceosome footprint as a single average. Recent developments in high-throughput branchpoint detection and machine-learning models (Signal et al. 2018) allowed us to identify branchpoints for the 81 3′SS in our data set. Branchpoint structure is likely to play a particularly important role in splicing regulation as branchpoint detection is the first step in defining the 3′ acceptor for spliceosome assembly (Reed and Maniatis 1988). We identified the 3′SS associated with each branchpoint allowing us to evaluate the strength of that splice site using SpliceAI (Jaganathan et al. 2019).

We performed k-means clustering (Shutaywi and Kachouie 2021) of RADIS-informed base-pair probabilities for the 42 nucleotides comprising each branchpoint spliceosome footprint and obtained three clusters (Figure 4D, 4E). One cluster (cluster 2) is distinct in showing a high level of structure downstream of the branchpoint. By contrast, cluster 1 has low overall base-pairing probability, consistent with open structures, and cluster 0 is characterized by intermediate base-pairing. Analysis of the sequences and structures does not reveal an obvious motif in cluster 2 that explains the high pairing probability downstream of the branchpoint; however, there is a modest relationship between base pairing probability and mean GC content for each cluster (Figure 4F). We do observe statistically different SpliceAI acceptor probabilities across the three clusters (Figure 4G); because SpliceAI scores splice sites rather than branchpoints, this is the acceptor probability of the 3′SS downstream of each branchpoint. Cluster 1 has the highest mean acceptor probability while cluster 0 has the lowest. Acceptor strength therefore does not track the overall base-pairing level of the footprint, since the most open class (cluster 1) shows the strongest acceptors and the intermediate class (cluster 0) the weakest. RADIS data therefore support dissection of branchpoint structures and reveal a functionally important set of structural features surrounding the spliceosome branchpoint footprint. Specifically, RADIS shows that branchpoint regions are not uniformly unstructured but instead fall into distinct structural classes, a form of per-branchpoint structural heterogeneity that transcriptome-wide methods have not resolved.

### RADIS analysis of deep intronic structures

Human introns make up 88% of transcribed nucleotides in pre-mRNAs, of which splice site sequence and structure comprise a small fraction (Waldern et al. 2022). Intronic sequences contain diverse RNA binding protein sites, repetitive elements, and splicing regulatory elements, all of which can affect splicing outcome. RADIS uniquely provides chemical probing structural information across these regions.

As one example, RADIS probes structure on both the positive and negative strands, and we therefore asked how reverse complement sequences of repetitive elements fold in pre-mRNAs. We focused on the 111 sense and 122 antisense full-length transcribed Alu elements in our dataset. We considered only elements that fall within annotated human pre-mRNA transcripts, so an Alu transcribed from the BAC but not transcribed in cells was excluded; the antisense Alu elements analyzed here are therefore transcribed in their native genomic context. Antisense Alu elements are of particular interest because inverted sense–antisense Alu pairs form the double-stranded RNA that is a principal substrate for ADAR editing (Athanasiadis et al. 2004) and that can recruit double-stranded RNA helicases such as DHX9, whose loss triggers aberrant splicing and circular RNA formation from inverted Alu repeats (Aktas et al. 2017). We first examined whether sense and antisense Alu elements are structured. Analysis of the mean DMS-informed base-pairing probability of sense and antisense Alu elements shows that antisense Alu elements are significantly less structured than sense ones (Figure 5A). This result is consistent with an analogous analysis of mean DMS reactivities, which are higher for antisense Alu RNAs (Figure 5B), even though there is no significant difference in GC content between our Alu sequences (Figure 5C). These analyses emphasize that sense Alu motifs form more compact base-paired structures independent of any underlying difference in sequence composition. Nonetheless, these comparisons show substantial element-to-element variation, suggesting significant structural diversity within each Alu strand.

**Figure 5:**
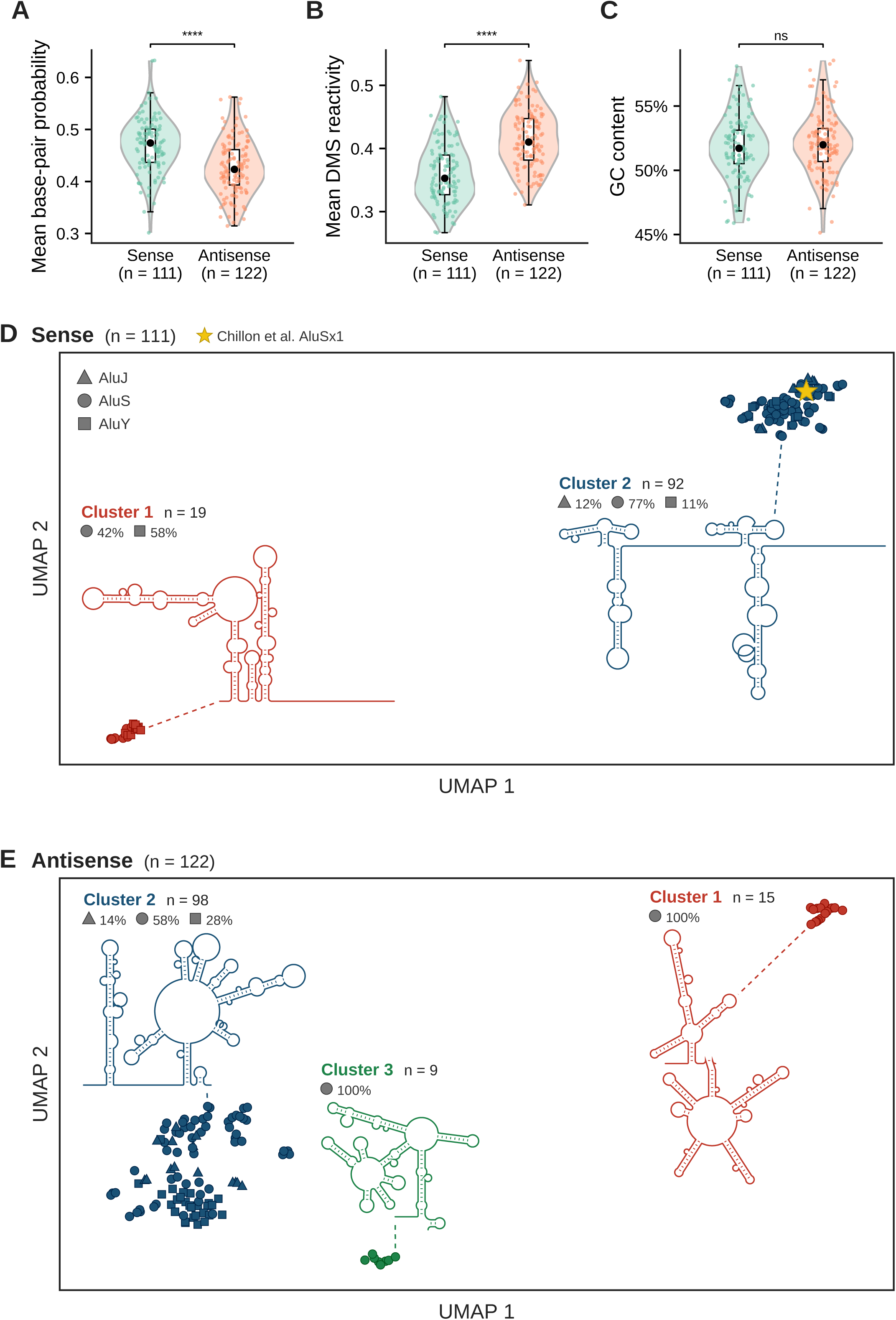
Secondary-structure landscape of sense and antisense Alu elements probed by RADIS. Per-element (A) mean DMS-informed base-pairing probability (B) mean DMS reactivity, and (C) GC content for full-length (>270 nt) sense (n = 111) and antisense (n = 122) Alu elements, shown as violin and overlayed box plots.****, p < 0.0001; two-sided Mann-Whitney U test. (D) Clustering of DMS-informed secondary structures for sense-strand Alu elements. UMAP plot shown embedding from a tree-distance matrix of mlocarna structure-based alignments. Structures partitioned into k = 2 clusters by k-medoids clustering of the tree-distance matrix. Cluster 1 (red, n = 19) and Cluster 2 (dark blue, n = 92) and are illustrated as representative R2R (Weinberg and Breaker 2011) covariation skeleton diagrams. Marker shapes denote Alu lineage (▴ AluJ, ● AluS, ▪ AluY) and AluJ/AluS/AluY lineage composition of each cluster is shown. (E) Clustering analysis for antisense Alu elements. RNAs partitioned into k = 3 clusters: Cluster 1 (red, n = 15), Cluster 2 (dark blue, n = 98) and Cluster 3 (green, n = 9).

We next asked how similar the folds of individual Alu elements are to one another, computing a structure-based distance between every pair of elements on each strand. We projected the distance matrix into two dimensions with UMAP (McInnes et al. 2018). We partitioned the structures by k-medoids clustering of the tree-distance matrix, determining k using Silhouette Analysis (Shutaywi and Kachouie 2021), and used the embedding for display. Sense Alu RNA structures separate cleanly into two clusters (Figure 5D): a small cluster (n = 19) enriched in young AluY elements (58%, compared with 11% in the larger cluster), whose representative covariation structure folds the two Alu arms into a single continuous base-paired domain (Supplementary Figure 6A). The larger cluster (n = 92) is dominated by AluS-family elements (77%) whose representative covariation structure shows an open fold containing features consistent with the folding of ancestral 7SL (Zwieb 1985). Antisense Alu elements, by contrast, resolve into three clusters (Figure 5E; n = 15, 98 and 9), each with a distinct representative structure. Notably, most clusters contain a mixture of AluJ, AluS and AluY elements, so structural clustering does not simply recapitulate Alu phylogeny; the exceptions are antisense clusters 1 and 3, which are composed entirely of AluS elements (R2R (Weinberg and Breaker 2011) covariation models shown in Supplementary Figure 6B).

In sum, for both sense and anti-sense Alu elements, the local fold in its pre-mRNA context is influenced by both strand orientation and Alu sub-family. Importantly, RADIS data expose strand-dependent structural heterogeneity within a single class of repetitive element, uniquely obtainable by simultaneous probing of both strands of pre-mRNA.

## Discussion

RADIS decouples pre-mRNA structural interrogation from cellular transcription and degradation by transcribing human genomic loci from plasmid (BAC) templates and probing the resulting RNA by DMS-MaP (Krokhotin et al. 2017; Mustoe et al. 2019; Grayeski et al. 2022). Three innovations make RADIS suited to long RNAs such as pre-mRNAs: (*i*) library construction starts from 150– 170 kb of human genomic sequence rather than cellular mRNA, removing the abundance penalty that has limited transcriptome-wide intronic coverage (Ding et al. 2014; Rouskin et al. 2014; Bubenik et al. 2021; Eich et al. 2024; Yarshova et al. 2026); (*ii*) a modified hairpin adaptor (NEBNext) appends the T7 promoter, enabling *in vitro* transcription of high-complexity, strand-resolved RNA pools without per-locus primer design; and (*iii*) a directional cDNA workflow preserves strand resolution, so sense and antisense reactivities can be analyzed separately. Any sequence accessible via a large-scale DNA template is amenable to RADIS; here we probed 109 5′SS, 88 3′SS and 233 full-length Alu elements, an expansion of up to two orders of magnitude over prior targeted pre-mRNA probing.

Two complementary benchmarks validate RADIS reactivity data. First, for *E. coli* ribosomal RNAs (obtained via chromosomal DNA carry-through), RADIS reactivities correlate strongly with cell-free DMS-MaP data (Pearson r = 0.81–0.87), and DMS-constrained modeling recovers known 16S and 23S architectures with AUC values of 0.74–0.75, closely matching the cell-free reference (0.78–0.79) (Hajdin et al. 2013). Second, at seven splice sites, RADIS reactivities agree closely with in-cell DMS probing (r = 0.87–0.90). *BRCA1* is Alu-rich, and a substantial fraction of pathogenic alleles arises through Alu-mediated deletions or cryptic exonization (Mazoyer 2005; Yu et al. 2026); the per-element structural classification reported here creates a reference against which allele-specific structural perturbations can be benchmarked. For the *CRB2*–*DENND1A* locus, the alternatively spliced *DENND1A*.V2 isoform, which is causally linked to polycystic ovary syndrome (McAllister et al. 2014; Tee et al. 2016), is expressed at low levels in disease-relevant theca cells. RADIS could therefore provide an experimentally supported scaffold complementing sequence-based predictions (Halvorsen et al. 2010; Bonnal et al. 2020).

Our study includes several limitations. RADIS probes RNA outside of cells. In-cell folding can be modulated by RNA-binding proteins, chromatin context, and co-transcriptional kinetics that are absent in our cell-free transcription environment (Saldi et al. 2021). Because tertiary contacts in folded RNAs tend to span fewer than ∼200 nt, most three-dimensional structure is likely captured; interactions spanning more than ∼300 nt, however, are underrepresented as a consequence of the ∼1.5 kb fragmentation of DNA template. Longer fragment distributions and, potentially, more processive enzymes could extend the accessible interaction range.

Several extensions are immediately tractable. First, the strand-resolved probing readout is ideally suited to systematic mapping of antisense transcripts and overlapping convergent transcription units, which transcriptome-wide methods do not resolve. Second, the workflow is compatible with any probing chemistry. Ultimately, RADIS is a generalizable framework for converting a plasmid-resident genomic locus into a quantitative RNA structural model, providing the missing link between sequence-based predictions and probing-informed RNA structure models and their effects on biological regulatory mechanisms.

## Materials and Methods

### Cell culture

T-47D cells (ATCC number HTB-20) were cultured in ATCC-formulated RPMI-1640 Medium (ATCC #30-2001) supplemented with human recombinant insulin (Gibco #12585-014; 1.0 mL per liter), 10% fetal bovine serum (ATCC #30-2020) and 0.5% pen/strep (Gibco). Cells were maintained at 37°C and 5% CO₂.

### BAC culture and extraction

BAC clones RP11-95G8 (*ESR1*), RP11-147E3 (*CRB2*–*DENND1A*) and RP11-812O5 (*BRCA1*), in the pBACe3.6 vector and maintained in *E. coli* DH10B, were obtained from BACPAC Genomics (Berkeley, CA, USA). BAC-containing bacteria were grown on LB-agar plates supplemented with 12.5 µg/ml chloramphenicol and single colonies were used to inoculate liquid LB cultures with 12.5 µg/ml chloramphenicol. Overnight cultures were grown at 30°C with shaking, and BAC DNA was purified (NucleoBond Xtra Midi kit; Macherey-Nagel, cat. no. 740410.50). Purified BAC DNA was quantified (Qubit fluorometer; Thermo Fisher Scientific) and used directly for fragmentation and library preparation. The integrity of the BAC preparations was verified by pulsed-field electrophoresis on a PFGE system (Bio-Rad).

### RNA preparation from BAC DNA

BAC DNA (500 ng) was diluted to 130 µL in Tris-EDTA (TE) buffer and sheared (Covaris acoustic sonication; 1% intensity, 20 seconds) to generate fragments above 1000 bp. Size selection was performed: dual-ratio AMPure XP bead purification: beads were first added at 0.45× ratio to capture large fragments, and the supernatant was then cleaned at 0.55× ratio to recover the desired size range. DNA was eluted in 20 µL of 0.1× TE buffer. Fragment size and concentration were verified by TapeStation analysis and Qubit fluorometry, respectively. Libraries were constructed (NEBNext Ultra II DNA Library Prep Kit; New England Biolabs) and between 200– 500 ng of size-selected DNA was subjected to end repair and dA-tailing, followed by adaptor ligation (1:5 dilution). PCR amplification was carried out for 6–8 cycles using custom primers: Adaptor 1 with T7 promoter (5’-TAATA CGACT CACTA TAGGA CTGGA GTTCA GACGT GTGCT CTTCC GATC-3’) and Adaptor 2 (5’-CACTC TTTCC CTACA CGACG CTCTT CCGAT C-3’).

Libraries were purified (AMPure XP beads, 0.8× ratio), eluted in 20 µL TE buffer, and verified by TapeStation and Qubit analysis. RNA was synthesized in vitro (HiScribe T7 High Yield RNA Synthesis Kit; New England Biolabs) from 150 ng DNA template at 37°C for 2 hours and purified (NEB RNA Cleanup Columns), yielding several µg. The RNA was treated with Turbo DNase (1 µL in 50 µL reaction, 37°C, 15 minutes; repeated once) and purified again using NEB RNA Cleanup Columns. RNA concentration was quantified by NanoDrop spectrophotometry, and integrity was verified by TapeStation analysis.

### Chemical probing of RNA *in vitro*

RNA (3 µg) was diluted in 15 µL molecular biology grade water and heat denatured at 98°C for 1 minute, followed by snap-cooling on ice for 1-2 minutes. Denatured RNA was combined with 15 µL of 2× Bicine folding buffer (600 mM Bicine pH 8.3, 200 mM NaCl, 10 mM MgCl₂) (Mustoe et al. 2019) and refolded at 37°C for 30 minutes. Nine microliters of folded RNA was then added to either 1 µL of 1.7 M DMS (final concentration 170 mM) prepared in a 1:2 (v/v) nitromethane/sulfolane (NS) solution or 1 µL of NS solution as a vehicle control. Reactions were incubated at 37°C for 6 minutes before quenching with an equal volume of 1:3 (v/v) β-mercaptoethanol (BME) in water. RNA was buffer exchanged (Sephadex G50 spin column) prior to downstream analysis.

Probed RNA (8.8 µL) was combined with 2 µL of 10 mM dNTPs (NEB) and 1 µL of 2 µM adaptor 2 primer. This mixture was heat denatured at 98°C for 1 minute and then chilled on ice for 1-2 minutes. Added to each reaction was 2 µL of freshly prepared 10× NTP minus first strand buffer (0.5 M Tris pH 8.0, 0.75 M KCl, 0.1 M DTT), 4 µL of 5 M betaine, and 1.2 µL of 100 mM MnCl₂. Reactions were incubated at 25°C for 2 minutes before adding 1 µL of either SuperScript II RT enzyme or water (for no-RT controls). Reverse transcription was performed with the following program: 25°C for 10 minutes, 42°C for 90 minutes, 10 cycles of 50°C for 2 minutes followed by 42°C for 2 minutes, then 70°C for 10 minutes, and a hold at 12°C. cDNA was buffer exchanged with Sephadex G50 spin columns prior to second strand synthesis.

The second strand was synthesized (NEBNext Ultra II Directional RNA Second Strand Synthesis Module; NEB #E7550S). Sequencing libraries were prepared (NEBNext Ultra II FS DNA Library Prep Kit for Illumina; NEB), following the manufacturer’s protocol. PCR amplification was performed for 6 cycles, and libraries were purified using AMPure XP beads at a 0.8× ratio. Final library quality and size distribution were assessed by TapeStation analysis, and concentration was quantified by Qubit fluorometry. Libraries were pooled at equimolar concentrations and sequenced with the Illumina NextSeq 1000 platform.

### In-cell chemical probing

T-47D cells (300,000 per well) were seeded in 6-well plates and cultured for 48 h. Cells were harvested, washed with phosphate-buffered saline (PBS), briefly exposed to TrypLE, and incubated for 5 min at 37°C before resuspension in 1.0 mL bicine-buffered medium and transfer to 1.7-mL tubes maintained at 37°C. At least three biological replicates were generated from independent cell cultures separated by at least two passages.

For each sample, two 1.7-mL tubes were pre-warmed to 37°C containing 50 µL and 40 µL ethanol, respectively; immediately prior to probing, 10 µL DMS was added to the 40 µL aliquot. Cell suspensions (450 µL) were mixed and added to DMS or vehicle control tubes and incubated at 37°C for 6 min. Reactions were quenched with 0.5 mL of 25% β-mercaptoethanol (BME) and placed on ice. Cells were pelleted at 1000 ×g for 2 min at 4°C, the supernatant was removed, and pellets were lysed in 500 µL TRIzol and 100 µL chloroform followed by centrifugation at 12,000 ×g for 15 min at 4°C. The aqueous phase was recovered, and RNA was isolated by glycogen-assisted ethanol precipitation, washed with 70% ethanol, air-dried, and resuspended in 44 µL nuclease-free water.

RNA was treated with TURBO DNase and purified (NEB RNA Cleanup Kit), and 1 µg was subjected to MaP-RT as described for *in vitro* chemical probing, using primers specific for the splice sites analyzed (Supplementary Table 3). Reverse-transcription products were cleaned with 1.8× AMPure XP beads and amplified by two rounds of PCR with adaptor-containing primers. Libraries were pooled at equimolar concentrations and sequenced on an Illumina NextSeq 1000 instrument.

### DMS-MaP data processing and reactivity calculation

In-cell structure probing was performed in at least biological triplicate for each splice site. For analysis, replicate libraries were first processed separately with ShapeMapper2 (v2.1.5) in --dms mode (Busan and Weeks 2018) using the appropriate primer files, and replicates showing consistent reactivity profiles were concatenated to increase coverage. The combined in-cell datasets were then re-processed with ShapeMapper2 to obtain mutation-based DMS reactivity profiles, followed by visualization and downstream analyses with RNAvigate (v1.0.0) (Irving and Weeks 2024). DMS profiles, skyline plots and correlation plots were generated in RNAvigate using boxplot normalization, and correlation coefficients were calculated within RNAvigate.

For RADIS (*in vitro*) experiments, paired-end reads were first aligned to the appropriate BAC or *E. coli* rRNA reference using a strand-aware Bowtie2 (Langmead and Salzberg 2012) workflow in paired-end “forward–reverse” mode with local alignment and adjusted penalty parameters (--fr--no-mixed --sensitive-local --mp 3,1 --rdg 5,1 --rfg 5,1 --dpad 30 --maxins 600 --ignore-quals). Alignments were converted to BAM, filtered to retain mapped pairs, and coordinate-sorted with samtools (Li et al. 2009). To recover strand-specific read sets, BAM files were partitioned by FLAG combinations corresponding to read1/read2 orientation on the reference. These strand-specific BAMs were name-sorted and converted back to compressed FASTQ files using bam-to-FASTQ conversion and parallel gzip compression, yielding separate forward and reverse read sets for each condition. Strand-specific FASTQ files were then processed with ShapeMapper2 in DMS-MaP mode, with strand-resolved modified and untreated libraries supplied as separate paired-end inputs. ShapeMapper2 was run with a minimum depth threshold of 300 reads per position, per-read mutation histograms, and a 10-nt random primer length to produce per-nucleotide mutation rates and normalized DMS reactivities for each strand and condition. These reactivities formed the input for downstream comparative analyses, correlation calculations, and EternaFold-constrained secondary structure prediction.

### Ribosomal RNA analysis

Published *E. coli* ribosome DMS-MaP data were downloaded from GEO (accession GSE225383). For RADIS rRNA datasets, strand-sorted FASTQ files corresponding to reads aligned to *E. coli* 16S or 23S rRNA (replicate 2 for *ESR1* and *CRB2*–*DENND1A* BACs, and replicates 1 and 2 for the *BRCA1* BAC) were concatenated and processed with ShapeMapper2 in --dms mode to obtain per-nucleotide mutation rates and normalized DMS reactivities. RADIS and cell-free rRNA profiles were then globally rescaled to match the dynamic range of the published experiment before secondary structure prediction. Windowed secondary structure modeling was performed with UltraFold v1.0.0 (https://github.com/gsstephenson/UltraFold), a Python reimplementation of the windowed SuperFold approach (Smola et al. 2015) that folds overlapping windows in parallel and merges the resulting base-pairing probabilities into a single whole-RNA model. UltraFold was run with its EternaFold engine, which treats DMS reactivities as learned evidence rather than as a pseudo-free-energy term, and with a folding window size of 1000 and a step size of 300. Structures were visualized in RNAvigate (v1.0.0), enabling direct comparison of reactivity-guided 16S and 23S rRNA structures between RADIS and cell-free conditions. Consistent with recent benchmarks (Wayment-Steele et al. 2022), EternaFold constrained with DMS probing data showed good agreement with known rRNA secondary structures, and was therefore used for DMS-constrained folding analyses.

### Splice-site and branchpoint definition

Splice-site annotations for the BAC regions were obtained from the HEXEvent database by intersecting HEXEvent coordinates with each BAC insert. Alternative, constitutive and cryptic events were retained, yielding 109 5′ splice sites (5′SS) and 88 3′ splice sites (3′SS). To expand the set of cryptic splice sites, 18 publicly available RNA-seq datasets from untreated T-47D cells (SRA accessions SRR18944091–SRR18944093, SRR18889003–SRR18889005, SRR20614686–SRR20614688, SRR10919047–SRR10919052 and SRR19907131– SRR19907133) were downloaded, quality-filtered with Trimmomatic (Bolger et al. 2014), FastQC and BBDuk, and aligned to the human genome (hg38) using STAR with default parameters. The resulting BAM files were analyzed with MAJIQ (Vaquero-Garcia et al. 2016) to identify alternative and cryptic splice junctions, and MAJIQ-supported cryptic splice sites overlapping BAC regions were added to the HEXEvent set. Branchpoints were assigned for 81 introns based on published branchpoint maps (Mercer et al. 2015; Pineda and Bradley 2018; Briese et al. 2019; Zeng et al. 2022; Damianov et al. 2024) and predictions from the Branchpointer algorithm; when multiple candidate branchpoints were present, the position most frequently supported across datasets, or, when equally supported, the candidate closest to the 3′ splice junction was selected. For each splice site, a 42-nucleotide structural “footprint” was defined: 12 nt within the exon and 30 nt within the intron for 5′SS and 3′SS, and a 42-nucleotide window centered on the annotated branchpoint for branchpoint sites.

### Structure prediction and pairing scores

For splice-site analyses, a 400 nt sequence-window centered on each footprint was folded with EternaFold using RADIS DMS reactivities from the replicate with the most reads as constraints. For Alu and other repeat elements, 1000 nt windows-centered on the repeat were folded under the same conditions. Nucleotide-wise base-pairing probabilities were obtained from EternaFold Predict, and per-position pairing scores were computed as the sum of pairing probabilities for each nucleotide across all possible partners. For each footprint or repeat, the mean base-pair probability was calculated by averaging these perposition scores across the 42-nucleotide splice-site footprint or the full repeat window. Where indicated, ROC curves and AUC values were computed with RNAvigate by comparing predicted base pairs to accepted reference structures. SpliceAI (Jaganathan et al. 2019) scores were computed for all BAC sequences using default parameters, and SpliceAI scores corresponding to annotated splice junctions and cryptic sites were extracted for comparative analyses.

Differences in average base-pair probability among splice-site types (5′SS, 3′SS, and branchpoint) were assessed using non-parametric tests. Average base-pair probability was computed per site as the mean across 42 footprint positions, followed by an overall Kruskal– Wallis test and two-sided Mann–Whitney U tests for pairwise comparisons, with statistical significance defined as *p* < 0.05.

### Alu structural analysis

Full-length Alu elements (length > 270 nt) annotated within the probed BAC inserts and located within annotated human pre-mRNA transcripts were retained for structural analysis, yielding 111 sense and 122 antisense elements. For each element, the RADIS-derived per-nucleotide DMS reactivities were converted to soft constraints on base-pairing probabilities using the method of Zarringhalam et al. (Zarringhalam et al. 2012), yielding a per-position base-pairing probability profile that informed all downstream analyses. Per-element summary statistics (mean base-pairing probability, mean DMS reactivity, and GC content) were compared between sense and antisense pools using two-sided Mann–Whitney U tests (scipy.stats.mannwhitneyu). To examine structural heterogeneity within each pool, structure-based multiple sequence alignments were computed for the sense and antisense Alu elements separately with mlocarna, with the DMS-informed base-pair probabilities supplied as RNA dotplot files so that the alignment was biased toward chemical-probing-supported pairings (Zarringhalam et al. 2012). mlocarna uses the SPARSE implementation in LocARNA (Local RNA Alignment), which leverages base-pairing probabilities, to accelerate the Sankoff algorithm (Will et al. 2012; Will et al. 2013; Will et al. 2015). The resulting tree-distance matrices from mlocarna were used as a precomputed metric for two-dimensional UMAP embedding (umap-learn 0.5.12), and cluster assignments were obtained by k-medoids clustering of the precomputed tree-distance matrix (k = 2 for sense, k = 3 for antisense), with the UMAP embedding used for display only and not to define clusters. For each cluster, a representative consensus structure was visualized as an R2R covariation diagram. Alu sub-family assignments (AluJ, AluS, AluY) were taken from RepeatMasker labels, and per-cluster lineage composition is shown as bar-chart insets (Figures 5D and 5E).

## Acknowledgements

This work was supported by U.S. National Institutes of Health grants (R01 HL111527 to A.L. and K.M.W. and R35 GM140844 to A.L., and R01 HL124233, R01 HL166992, and R01 HL171213 to P.J.C.).

## Supporting Information

Sequencing data are accessible through Gene Expression Omnibus (GEO) accession number GSE341315, accessed from NCBI. Analysis code and processing pipelines are available at https://github.com/ClaireFleur/RADIS and https://github.com/gsstephenson/UltraFold.

